# LR-DNase: Predicting TF binding from DNase-seq data

**DOI:** 10.1101/082594

**Authors:** Arjan van der Velde, Michael Purcaro, William Stafford Noble, Zhiping Weng

## Abstract

Transcription factors play a key role in the regulation of gene expression. Hypersensitivity to DNase I cleavage has long been used to gauge the accessibility of genomic DNA for transcription factor binding and as an indicator of regulatory genomic locations. An increasing amount of ChIP-seq data on a large number of TFs is being generated, mostly in a small number of cell types. DNase-seq data are being produced for hundreds of cell types. We aimed to develop a computational method that could combine ChIP-seq and DNase-seq data to predict TF binding sites in a wide variety of cell types. We trained and tested a logistic regression model, called LR-DNase, to predict binding sites for a specific TF using seven features derived from DNase-seq and genomic sequence. We calculated the area under the precision-recall curve at a false discovery rate cutoff of 0.5 for the LR-DNase model, a number of logistic regression models with fewer features, and several existing state-of-the-art TF binding prediction methods. The LR-DNase model outperformed existing unsupervised and supervised methods. Additionally, for many TFs, a model that uses only two features, DNase-seq reads and motif score, was sufficient to match the performance of the best existing methods.

## INTRODUCTION

Transcription factors (TFs) play a fundamental role in the regulation of gene expression in the cell. The human genome encodes roughly 1,500 TFs which exert their regulatory functions by combinatorial binding to the genomic DNA via interaction with other regulatory proteins, leading to complex regulatory networks. In order to understand gene regulation, it is necessary to determine the genomic locations bound by each TF in specific cell types at specific developmental stages under specific conditions. Currently, chromatin immunoprecipitation followed by high-throughput sequencing (ChIP-seq) is the leading experimental technique for determining where a TF binds to genomic DNA in living cells. In an effort to annotate functional elements in the human genome, the ENCODE project (<u>http://encodeproject.org</u>) has performed a large number of ChIP-seq experiments on many TFs across a large number of cell types (1). However, the ChIP-seq assay can only profile one TF per experiment; hence, it is infeasible to use ChIP-seq to assay all TFs in all cell types under all conditions. Also, ChIP-seq experiments depend on the availability of ChIP-grade antibodies that are highly specific to the TF of interest, which poses a major challenge for many TFs.

DNase-seq is another experimental technique that can be used to probe regions of chromatin that are accessible to TF binding. In this technique, chromatin is first digested using the deoxyribonuclease I enzyme (DNase) which creates single-stranded nicks in the DNA, and deep sequencing is performed on short (~60 bp) fragments of DNA that are released by two DNase cuts (2). DNase has long been used to discover regulatory regions in open chromatin, called DNase I hypersensitive sites (DHSs), marked by increased sensitivity to cleavage by the enzyme (3–7).

Whereas DHSs are typically 200–250 base pairs (bp) in length, ultradeep DNase-seq data (hundreds of millions of reads for human) enable the detection of smaller sites (6–22 bp) within DHSs that are protected from DNase cleavage by proteins such as TFs that bind to these sites. The DNase cleavage profiles around TF protected sites, i.e., the number of DNase-seq reads whose 5´ ends lie at each position, are called footprints, and are characterized by a substantial depletion of DNase I cuts at bound sites relative to their flanking regions. Compared with ChIP-seq, the advantage of DNase-seq is that one DNase-seq experiment can potentially provide information on the sites bound by all TFs bound to the DNA. Hesselberth et al. used ultradeep DNase-seq (which they called “digital genomic footprinting”) to survey the binding sites of many TFs in the yeast genome (8). This assay was subsequently performed on human cells (9). The shapes of the resulting cleavage profiles are highly TF specific, likely caused by the local shape of the DNA while bound by the TF. Later studies indicate that the cutting bias of the DNase enzyme also contributes to the footprints; however, attempts to correct for this bias by incorporating a k-mer model of the DNase cleavage profile of chromatin-free DNA did not improve the ability of the footprints to predict TF binding sites (10).

A number of computational methods have been developed to predict TF binding sites using DNase-seq data. Algorithms such as Wellington, HINT and DNase2TF (11–13) scan DNase-seq data at every position in a genomic region and predict de novo sites that may be bound by TFs, agnostic to which TFs may bind there. Other methods take advantage of the binding motifs of TFs, models of which are accumulating rapidly, aided by high-throughput techniques like ChIP-seq, HT-SELEX (14), and protein binding microarrays (15). Cuellar-Partida et al. extended the FIMO algorithm, which was originally developed to scan a genome for motif sites of a TF given a position weight matrix, to incorporate a prior computed using DNase-seq data (16, 17). CENTIPEDE and PIQ (18, 19) first identify potential binding sites using the annotated binding motif of a specific TF, and then predict whether these motif sites are bound by the TF.

Computational methods can also be classified by whether they employ supervised or unsupervised learning techniques. Unsupervised methods do not rely on a training set in which the bound and unbound motif sites of a TF are known. CENTIPEDE (18) is an unsupervised method that uses an expectation maximization (EM) algorithm to fit a hierarchical model to classify motif sites into bound and unbound classes. The model incorporates features such as motif strength, cleavage profile, DNase-seq read counts, levels of histone modifications, evolutionary conservation, and distance to the transcription start site (TSS). PIQ (19) is another unsupervised method that models DNase cleavage profiles at nucleotide resolution. PIQ leverages multiple types of data as well as replicates, in order to make the method more robust to low coverage DNase-seq data. FIMO with epigenetic priors (17) is also an unsupervised method. In contrast, supervised methods are trained (i.e, their parameters are estimated) using a training set with known examples. BinDNase (20) is a generalization of the MILLIPEDE method (21). Both are supervised methods that use logistic regression and feature selection with features based on variable sized genomic regions.

To evaluate the performance of TF binding prediction methods, one requires a benchmark dataset containing known bound and unbound sites. This dataset can be constructed using ChIP-seq data of the respective TF, with candidate sites labeled “bound” or “unbound” based on whether they lie inside or outside of ChIP-seq peaks (genomic regions with statistically enriched ChIP-seq signal) for the TF. Because the resolution of ChIP-seq data are at the peak level (~250 bps) while binding sites are at the base-pair level, assumptions must be made while creating a benchmark dataset from ChIP-seq data. There are two widely used approaches, *site-centric* or *peak-centric*, to converting ChIP-seq data to bound and unbound sites (17). The site-centric approach concerns itself with only the motif sites for which the DNA sequence matches a given motif model with a score greater than a predefined cutoff. These sites are labeled “bound” if they lie within a ChIP-seq peak and “unbound” otherwise. All other positions in the genome, i.e., all sites that do not match well to the given motif model, are ignored. Note that, in the site-centric approach, multiple sites within a single ChIP-seq peak might pass the cutoff. In this case, each such site would be labeled “bound” by the site-centric approach. In comparison, the peak-centric approach labels the single best scoring motif site within each ChIP-seq peak as “bound” regardless of its score, and labels all other genomic positions as “unbound.” It is difficult to determine which approach is more biologically relevant. In practice, multiple sites in the same ChIP-seq peak may be bound either simultaneously or sequentially; thus, the site-centric approach would be correct in assigning all these sites as “bound.” On the other hand, when a ChIP-seq peak does not have any site above a motif cutoff, the best-scoring site may still be bound, in which case the peak-centric approach would be correct. Alternatively, an observed ChIP-seq peak may arise due to co-binding or indirect binding via interaction with another TF (22), in which case the site-centric approach would be correct in assigning no bound sites. In this paper, we focus on site-centric evaluation, which was also used for evaluating CENTIPEDE (18), PIQ (19), and BinDNase (20).

ChIP-seq data on a large number of TFs are being generated rapidly, primarily in a small number of cell types (1, 23, 24). Meanwhile, DNase-seq data are being produced for hundreds of cell types (25, 26), although many of these DNase-seq data sets are not at the ultradeep sequencing depth required for base-pair resolution characterization of DNase footprints. We aimed to develop a supervised computational method that could combine ChIP-seq and DNase-seq data to predict TF binding sites in a wide variety of cell types. We used logistic regression models to combine a standard motif score with biologically driven features of DNase cleavage signal around motif sites. We call our method “logistic regression of DNase” (LR-DNase). We trained LR-DNase using DNase-seq and ChIP-seq data in one cell type and made predictions using just the DNase-seq data in another cell type. Tested on five cell types with ChIP-seq data for 87 TFs, LR-DNase outperforms state-of-the-art methods that use high-resolution DNase cleavage profiles. We also show that a model that consists of just DNase read count and a motif score in many cases outperforms existing methods.

## METHODS

### DNase-seq and TF ChIP-seq data sets

We trained and tested our models on five cell types, K562, HepG2, GM12878, H1-hESC and HeLa-S3. Out of 371 ChIP-seq datasets on 88 TFs produced by the ENCODE consortium for these five cell types (<u>**Supplemental Table 1**</u>), a non-redundant set of 243 data sets was selected. ENCODE did not have a uniform processing pipeline for DNase-seq data, thus we processed these datasets as described below. On the other hand, ENCODE has a well-established pipeline for uniformly processing ChIP-seq data, thus we downloaded ChIP-seq peaks (genomic regions that are statistically enriched in the ChIP-seq signal) from the ENCODE portal (<u>https://www.encodeproject.org/</u>).

### DNase cleavage profiles

We mapped DNase-seq reads to the reference human genome (hg19) using the Bowtie algorithm (27) with the following setting: “bowtie --mm -n 3 -k 1 -m 1 -p 57 --chunkmbs 4000 -best --strata --phred64-quals --sam./local/bowtie/index/hg19 <fastq file>“. Reads were mapped to all chromosomes (including the mitochondrial sequence). ENCODE blacklisted regions and the entire mitochondrial sequence, which are known to accumulate spuriously large numbers of reads (1), were excluded for further analysis. We then extracted the DNase I cleavage signal by counting the 5´-ends of reads that mapped uniquely to one location of the genome and normalizing the total number of reads to 1 million. Each 5´-end represents a nick produced by the DNase I enzyme, located between its mapped position and the 3´ neighboring nucleotide.

### Annotation of TF binding sites

Previously we annotated the canonical sequence motif for the TF in each of the ENCODE ChIP-seq dataset and constructed the corresponding position-weight matrix (PWM) for the motif (22). In this study, we used the FIMO algorithm (16) to scan the entire human genome to find sites that match each motif with a p-value less than or equal to 10^−5^. These motif sites constitute the working space of all algorithms in this study. In other words, we assume that a TF would not bind to other locations in the genome. A p-value threshold of 10^−5^ is lenient, leading to 9,534– 446,680 motif sites (median 84,231) for the TFs in this study. In a particular cell type, the motif sites that fall within the ChIP-seq peaks—derived from the ChIP-seq experiment performed on the same TF in the same cell type—are considered as the binding sites of the TF in that cell type (positive examples). Conversely, the motif sites outside ChIP-seq peaks are considered not bound by the TF in that cell type (negative examples) (<u>**Supplemental Figure 14**</u>). We constructed such a gold standard for all cell types and used them for evaluating the algorithms in this study. Among the 243 TF and cell type combinations, 5.34±6.42% of the motif sites were labeled as positive examples. Thus these are highly unbalanced datasets, which are challenging for machine learning algorithms.

### LR-DNase

We developed the LR-DNase method, which models the log-likelihood-ratio that a motif site is bound by a TF based on the following logistic regression formula:

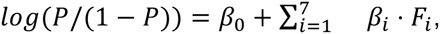

where P is the probability (or likelihood) that the site is bound by the TF, are the weights for each feature (*F_i_)*, and we considered seven features in total, detailed below. To make the scales of the features comparable, each feature was standardized to a Z-score. The model was trained with L1 regularization to prevent overfitting, using the glmnet (28) package in R. Nested cross-validation was performed, with three-fold cross-validation to evaluate the algorithm and ten-fold cross-validation within each of the three training sets to learn the regularization parameter lambda. In order to ensure inclusion of similar numbers of positive examples in each fold, we used a stratified cross-validation strategy in which training data for positive and negative examples were prepared separately.

### Features in LR-DNase

At each motif site, four features were extracted from DNase-seq data, complemented by three features derived from genomic sequence, gene annotation, and evolutionary conservation. In the region surrounding each motif site, we normalized the DNase cleavage signal by averaging cuts over positions with at least one read, in the 10,000-bp window centered at each position of the region. The normalized counts were log-transformed.

1. *DNase reads:* the total number of reads in a region of ±100 bp plus motif size centered at the motif site
2. *DHS*: a binary flag with value 1 or 0, indicating whether or not a motif site overlaps with a DNase I hypersensitive region (DHS). DHS regions were called using the HOMER software (29) with the following parameters: “findPeaks -region -size 500 -minDist 50 -o auto -tbp 0”. With these settings, HOMER detected regions with sizes of multiples of 500 bp separated by at least 50 bp.
3. *flank-to-footprint ratio*: the ratio of the average number of reads per nucleotide within the motif site (M) and the average number of reads per nucleotide in the left (L) and right (R) 100-bp regions flanking the motif, i.e., (M+1)/(L+1)+(M+1)/(R+1), where one read was added to prevent noise caused by division of two small values.
4. *strand difference score*: the difference between the total number of reads mapped to the Watson strand and the total number of reads mapped to the Crick strand. In both 100-bp regions flanking the motif site, the per-strand average number of reads per base was determined for each strands and the difference between the averages was taken and summed.
5. *motif score*: the log-likelihood ratio score for the motif site produced by the FIMO algorithm (16).
6. *evolutionary conservation:* PhastCons (30) conservation score at the motif site.
7. *distance to the nearest transcription start site*: the genomic distance of the motif site to the nearest transcription start site (TSS) of GENCODE (v19) genes (31).

### Within–cell-type and cross–cell-type predictions

For each TF/cell type combination, the positive and negative examples in the gold standard as defined above were divided separately into a training set, which contains two-thirds of the positive examples and two thirds of negative examples, and a test set, which contains the remaining one-third of the examples. Models were trained on each training set and then tested on the test set. DNase-seq data was available for all five cell types, but ChIP-seq data for most TFs was available for only some of the cell types. For each TF/cell type combination for which ChIP-seq data was available, the model was trained on each cell type and tested on all others.

### Logistic regression models using a subset of the seven features

We compared the performance of LR-DNase based on all seven features with a simpler logistic regression model containing two features, DNase reads and motif score. Additionally, we compared LR-DNase to each ranked feature separately.

### Comparison with other methods

We compared LR-DNase with four state-of-the-art methods, CENTIPEDE, PIQ, FIMO with epigenetic priors and BinDNase (17–20). Each method was run with default parameters and evaluated on the same set of motif sites for CENTIPEDE, BinDNase, and FIMO with a DNase-seq derived prior. PIQ performs its own motif search. To ensure that PIQ made predictions on all motif sites in our set, we used the maximum allowed number of motif sites where necessary.

### CENTIPEDE

CENTIPEDE was run with default parameters, DampLambda = 0, DampNegBin = 0, TrimP = 1e-04, NRiter = 5, and sweeps (max iterations) = 100.

### PIQ

PIQ performs its own motif search and cannot be used to make predictions on a custom set of motif sites. Accordingly, we increased the maximum number of motif site detections in cases where the number of motif sites in our data set exceeded the default for PIQ.

### BinDNase

The feature selection component of BinDNase is prohibitively slow when there are a large number of training examples. We ran BinDNase with training sets sized similar to what was used in the published paper. For within–cell-type training and testing, each dataset was divided into a ⅓ training and ⅔ test set. The training set was then subsampled to a maximum of 3000 positive and 3000 negative examples. The same models were used to perform cross–cell-type testing.

### Evaluation score

Because AUC-ROC normalizes out the skewness in the test dataset (i.e., the relative number of positive vs. negative labels), the performance of each method was measured by AUC-PR instead, at a FDR cutoff of 0.5. We calculate this AUC on a monotonized PR curve, where the precision at each threshold is replaced by the maximum precision at this threshold or any threshold further down the list.

### Fraction of maximally achievable gain in performance (FMI)

Fraction of maximum achievable improvement (FMI) over baseline model LR2 was calculated as (*m*-*b*)/(1-*b*) for *m* > *b* and (*m*-*b*)/*b* for *m* < *b*, where *b* is the AUC-PR50 for LR2 and *m* the AUC-PR50 for the model to be compared. The FMIs achieved by the best methods were 2.92±18.79%, median 2.95%.

## RESULTS

### The LR-DNase algorithm

Our LR-DNase algorithm uses L1 regularized logistic regression of seven features (described below, and see Methods for details) to model the log-likelihood-ratio that a motif site is bound by a TF. We model TF binding in a site-centric fashion, i.e., rather than making a prediction for each position in the genome, our method first scans the genome for potential binding sites using a position-weight matrix. Positions that contain a motif occurrence scoring above a pre-specified score threshold are then classified by LR-DNase as bound or not bound. In this respect, LR-DNase is similar to methods such as PIQ, CENTIPEDE and BinDNase.

The first four features of LR-DNase attempt to capture the most prominent commonalities of DNase cleavage profiles (henceforth abbreviated as “DNase profiles”) around the motif sites (or “sites” in short) that are bound by a TF, defined as the positions within ChIP-seq peaks of the TF that achieve position-weight matrix scores better than a specified threshold. <u>**Figure 1**</u> shows the average DNase profiles centered on the bound sites of three example TFs (CTCF, USF1 and NRF1). <u>**Supplemental Figure 1**</u> shows the DNase profiles for the bound sites of all TFs in this study and as negative controls, DNase profiles for the sites that are not bound by the TF as well as DNase profiles computed using chromatin-free and TF-free DNase-seq data (32). Although the DNase profiles are highly variable among the TF bound sites across the genome, they do show distinct characteristics from sites that are not bound. The first two features of LR-DNase (*DNase reads* and the binary *DHS flag*) are based on the observation that most TFs bind to open chromatin regions that are characterized by a large number of DNase reads (note the different ranges of the Y-axes for the sites that are bound and not bound by a TF in <u>**Supplemental Figure 1**</u>). The *flank-to-footprint ratio* feature quantifies the decrease in DNase reads at the bound sites due to protection by the bound TF, in comparison with the flanking regions, signified by the dip in the center of the DNase profiles at bound sites (<u>**Figure 1**</u> and <u>**Supplemental Figure 1**</u>). This feature is analogous to the footprint occupancy score proposed previously by Neph et al.(9). Furthermore, DNase cleavage profiles around TF bound sites show an imbalance between the cuts on the Watson and the Crick strands (blue and red curves in <u>**Figure 1**</u> and <u>**Supplemental Figure 1**</u>), which was reported earlier by Piper et al.(11). We use the *strand difference score* feature to quantify this imbalance.

**Figure 1.**
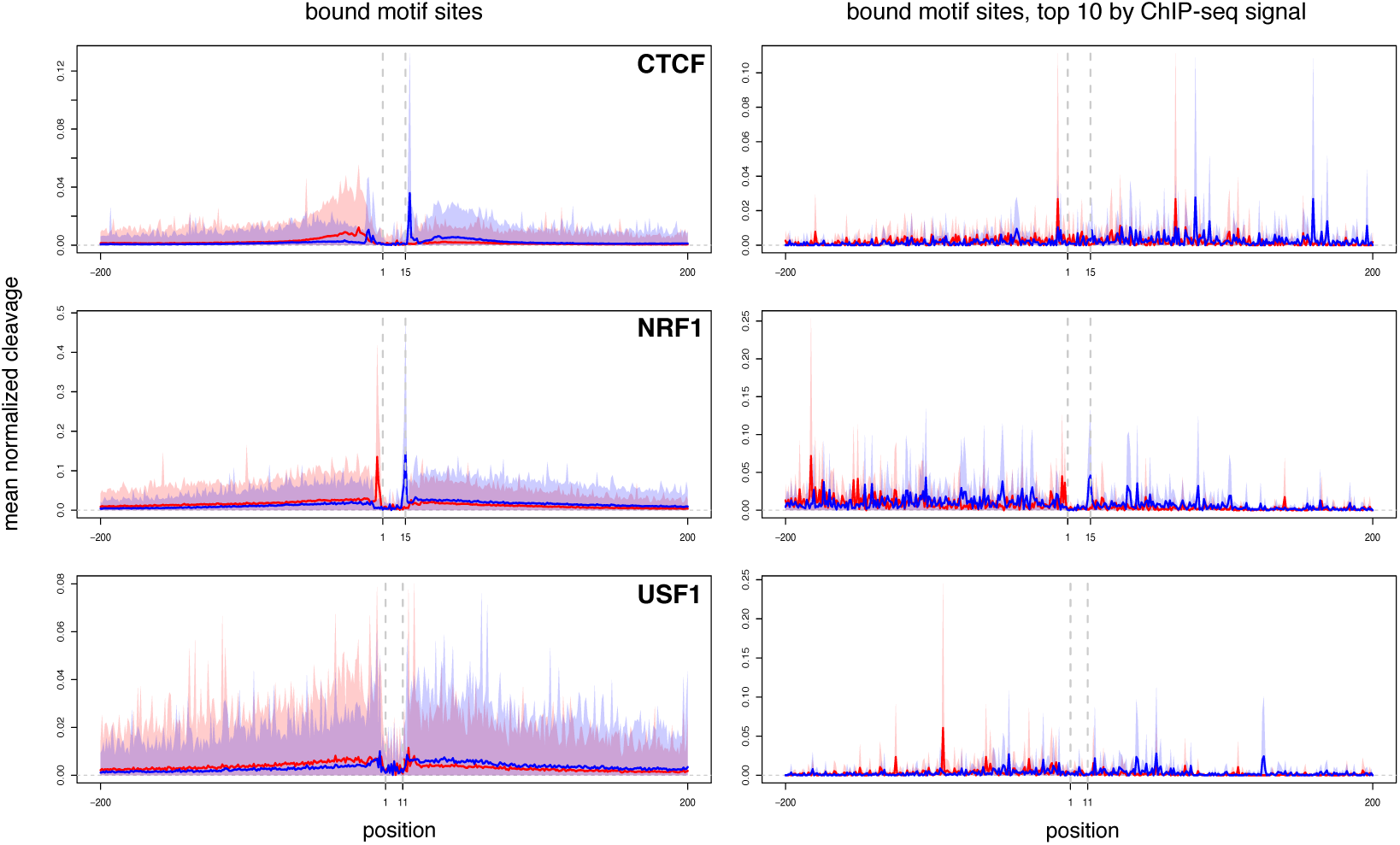
The average DNase cleavage profiles for CTCF, USF1 and NRF1 in K562 cells. The DNase profiles averaged over all bound motif sites in ChIP-seq peaks are shown in A-C. D-F show the average profile for the top 10 motif sites, ranked by ChIP-seq signal. The X-axis shows the genomic position relative to the motif site and Y-axis shows the mean of the normalized DNase cleavage counts. At every bound motif site, the DNase cuts were counted in a window flanking each motif site by 200 bps on both sides. The average number of cuts are shown as a solid red line for the reads mapped to the Watson strand and blue for reads mapped to the Crick strand. The standard deviations at each position (mean±sd) are shown as shaded red and blue areas. Dashed grey vertical lines mark the motif site. Each dataset was normalized to a total of 1 million reads.

*Motif score* is the fifth feature, computed by the FIMO algorithm (16) as a log-likelihood score that quantifies how well a site matches the annotated motif. Finally, we considered the average PhastCons (30) *evolutionary conservation score* at the motif site and the *distance of the motif site to the nearest transcription start site* (TSS).

### Evaluation of the performance of LR-DNase

We evaluated the performance of LR-DNase on five cell lines for which DNase-seq data as well as ChIP-seq data for many TFs were available from the ENCODE consortium (1): K562, HepG2, GM12878, HeLa-S3, and H1-hESC. For each TF, we first scanned the entire human genome using the annotated motif for the TF (22) and only retained sites that were more significant than an unadjusted p-value cutoff of 10^−5^, which is lenient for the ~6 × 10^9^ bp of genomic DNA (considering both genomic strands). Then in each cell type, we defined the motif sites that fell within the ChIP-seq peaks of the TF as bound by the TF, and the remaining as unbound. The fractions of sites that were designated as bound ranged from 0.01% to 26.30% (median 2.49%), as shown in <u>**Supplemental Table 1**</u>. The goal of LR-DNase is accurate classification of bound motif sites vs. unbound sites. We performed three-fold cross validation on the sites within each cell type, i.e., we trained LR-DNase using two-thirds of randomly chosen sites (with the percentage of bound sites preserved) and tested it on the remaining one-third of the sites, repeating this procedure three times, each with a different one-third used as the test set. We also trained LR-DNase using the data in one cell type and tested it on motif sites of the same TF in a different cell type (cross–cell-type performance).

The area under the curve (AUC) of the receiver operating characteristic (ROC) curve is often used to measure the performance of a binary classification algorithm; however, ROC analysis effectively normalizes the relative percentages of positives (bound motif sites) and negatives (unbound motif sites) to be 50% each. Because our datasets are highly skewed (the median percentage of positives is 2.49%), AUC-ROC is inappropriate. In contrast, the AUC of a precision-recall (PR) curve directly reflects the skewness of the dataset and is hence a more appropriate measure of algorithm performance on genomic datasets (33, 34). <u>**Figure 2**</u> shows the PR and ROC curves for the FOSL1 TF in H1-hESC cells. Among the 28,060 motif sites in the test set, only 16 were bound according to the ChIP-seq data of FOSL1 in this cell type. For such a skewed dataset, all LR-DNase models had very low precision (5-10%), yet they achieved nearly perfect AUC-ROC (0.99). The models could easily rank the 16 positives in the top 1% of the of the sites (hence AUC-ROC=0.99); nevertheless, over 90% of the top 1% sites, which amounted to 280 sites, were false positives (hence AUC-PR<10%). This example underscores the importance of using the AUC-PR for performance evaluation. The null performance for AUCROC is 0.5, and the null performance for AUC-PR is the fraction of positives in the test set (6 × 10^−4^ for this example). In other words, the task of identifying the positives is inherently more difficult if the fraction of positives is lower. A corollary is that the AUC-PR values for two TFs cannot be directly compared if they have different fractions of bound sites.

**Figure 2.**
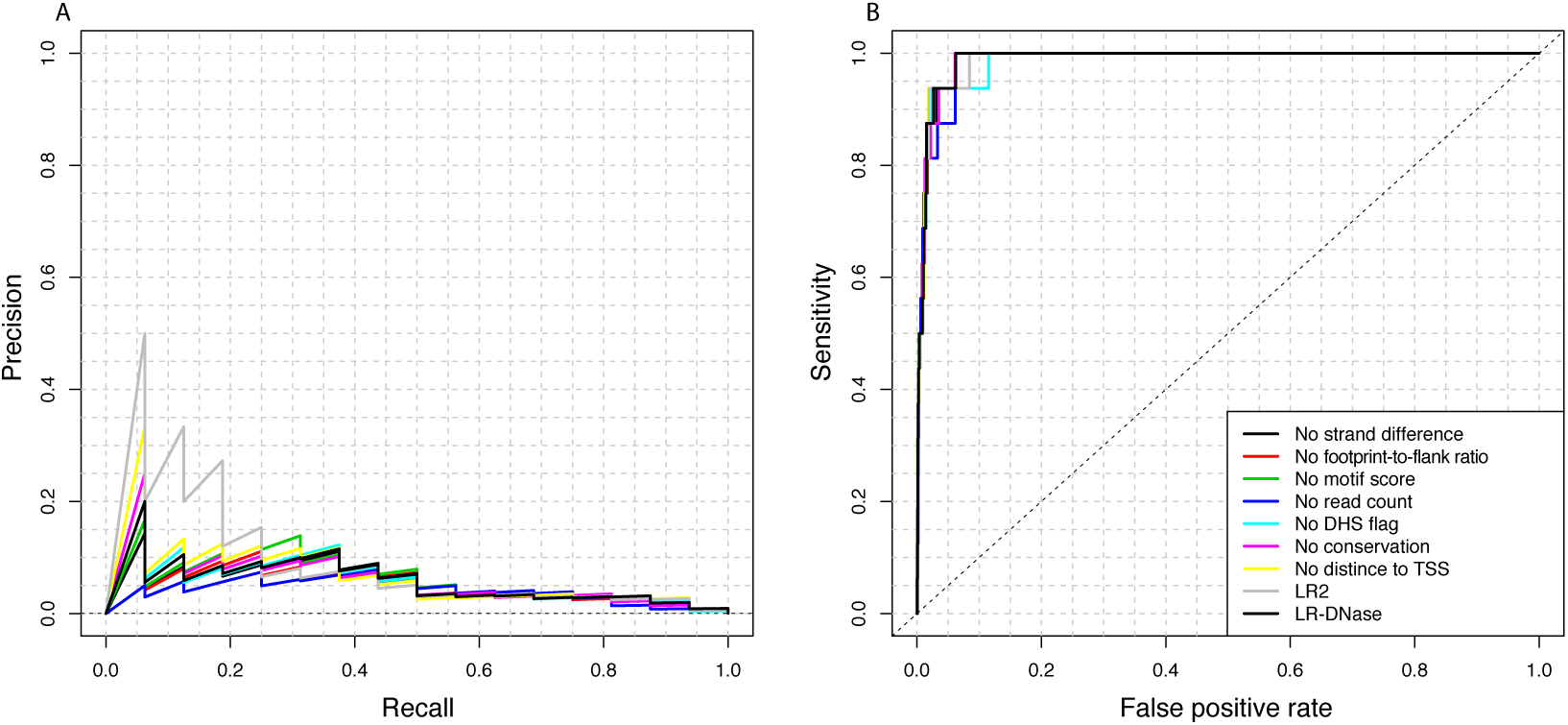
Precision-recall (PR) and receiver operating characteristic (ROC) curves for the transcription factor FOSL1 (AP-1 motif) based on within–cell-type cross validation in H1-hESC cells. The ROC curves for all LR-DNase models were nearly perfect (0.99±0.001), but the corresponding PR curves revealed low precision. Only 16 of the 28,060 motif sites in the test set were bound by AP-1, hence the jagged shape of the curve. Note that sensitivity is the same as recall.

When tested using three-fold cross validation, LR-DNase performed much better than the null model, i.e., the fraction of positives, for all datasets (<u>**Figure 3A**</u>). For datasets with ≥1% positives (180 out of 243), the AUC-PR for LR-DNase was 0.64±0.18 (mean±standard deviation). For datasets with fewer than 1% positives, LR-DNase performed with varying degrees of success, with AUC-PR at 0.31±0.21. The datasets with AUC-PR lower than 0.4 all had lower than 8.3% positives, i.e., it was difficult to identify the small numbers of positives.

**Figure 3.**
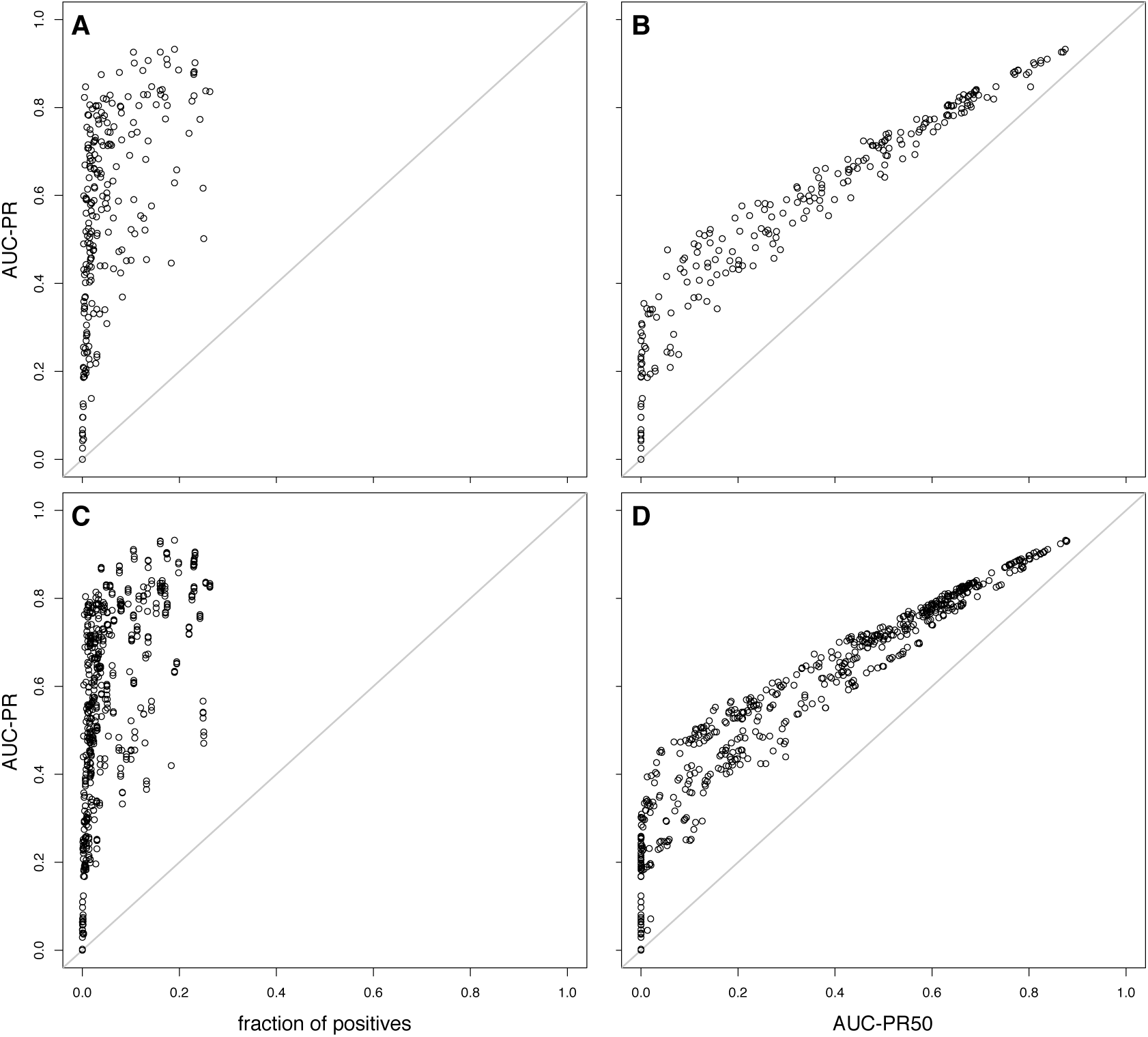
Comparison of LR-DNase with the null model using AUC-PR and AUC-PR50 as the performance measure, in the within–cell-type (panels A and B) and cross–cell-type (panels C and D) settings. (A) AUC-PR of LR-DNase as a function of the fraction of positives in the test set. The AUC-PR of the null model equals the fraction of positives in the test set, indicated by the grey diagonal line. For the datasets with the fractions of positives greater than 0.01, the AUC-PR was 0.64±0.18. (B) AUC-PR vs. AUC-PR50. AUC-PR50 is a stricter measurement of performance, emphasizing lower FDR. A number of datasets for which the AUC-PR was greater than 0 had AUC-PR50 of zero. (C) As in (A) but for cross– cell-type predictions. For the datasets with the fractions of positives greater than 0.01, the AUC-PR was 0.64±0.17. (D) As in (B) but for cross–cell-type predictions.

In a realistic situation one would pick a number of top predictions for experimental testing. A high number of false positives among the top predictions would therefore render the method inadequate. A false discovery rate (FDR) cutoff of 5% is commonly used to control the number of false positives. However, none of the state-of-the-art methods, including LR-DNase, yielded good results at such a stringent cutoff—for 50% of the datasets LR-DNase yielded an AUC-PR of zero at FDR thresholds of 5%. Nevertheless, in order to emphasize the requirement for a low FDR, we calculated the AUC-PR at FDR<0.5, or AUC-PR50 (<u>**Figure 3B**</u> and <u>**Supplemental Figure 2**</u>), and used it as the metric for comparing all the methods. Depending on the throughput of the follow-up experimental validation, FDR of 50% can still be highly informative in practice. Since none of the datasets has more than 50% positives, the null for AUC-PR50 is zero for all datasets. The AUC-PR50 of LR-DNase was greater than zero for 213 out of 243 datasets, covering 81 of the 87 TFs we studied.

We tested how well LR-DNase generalized across cell types by training a model on the DNase-seq data for a given TF in one cell type and testing the model with the DNase-seq data in a different cell type. We performed this analysis on all pairs among the five cell types (<u>**Figure 3CD**</u>). We observed substantial variations in the differences in performance between the within– cell-type prediction and the cross–cell-type prediction, especially for datasets with fewer than 1% positives. However, for the 180 datasets with ≥1% positives, cross–cell-type AUC-PR50 values were 91.16±19.71% of the within–cell-type performance (<u>**Supplemental Figure 3**</u>).

For several TFs LR-DNase performed substantially worse in the cross–cell-type setting than in the within–cell-type setting (labeled in <u>**Supplemental Figure 3**</u>), and NFE2 was the most extreme case. The ChIP-seq data were for two cell types, GM12878 and K562. The AUC-PR50 of the LR-DNase model trained on K562 was 0.50 for within–cell-type cross validation but only 0.01 for predicting GM12878 sites. There were only six bound sites in GM12878 (note that a bound site must pass the p-value threshold of 10^−5^ for matching the NFE2 motif), while the NFE2 ChIP-seq peaks in this cell type were highly enriched in the motif of another TF, USF1 (96% peaks contained an USF1 site), suggesting that the ChIP-seq peaks arose from the interaction between the NFE2 and USF transcription factors. NFE2 has been shown to interact with USF to cooperatively regulate the β-globin gene in erythroid cells (35). The NFE2 ChIP-seq peaks in K562 cells on the other hand, were highly enriched in the NFE2 motif (75.98% of the 1991 peaks matched the NFE2 motif) and less enriched in the USF1 motif (15.59% peaks matched the USF1 motif). Thus the LR-DNase model trained on the NFE2 sites in K562 would not be effective in predicting the few NFE2 sites in GM12878.

### Contributions of the features of LR-DNase

We took two complementary approaches to evaluating the contribution of each of the seven features of LR-DNase. First, we simply ranked the motif sites using each feature, i.e., no model was trained; rather, the ranks were used directly to plot the PR curve. <u>**Figure 4** **(top panel)**</u> shows the performance (measured in AUC-PR50) of each individual feature compared with that of the full LR-DNase model with all seven features. In all five cell types, *DNase reads* and *flank-to-footprint ratio* were the two most predictive features: their performance was 85.91% and 60.05% of that of the full model by median, respectively. The third most predictive feature was the *strand difference score* (median 28.94%). All these three features were computed from DNase-seq data. There is a moderate correlation (mean r=0.37±0.13) between *DNase reads* and *flank-to-footprint ratio,* and a weak negative correlation between *DNase reads* and *strand difference score* (mean r=–0.16±0.15) (**Table 1**).

**Table 1.** Spearman correlation coefficients between any two features of LR-DNase. Both mean and standard deviations are shown, computed using all non-redundant data sets (n=243).

|  | Flank-to-footprint ratio | Motif score | DNase reads | DHS flag | Dist. nearest TSS | Conervation |
| --- | --- | --- | --- | --- | --- | --- |
| Strand difference score | **-0.23 (0.13)** | -0.04 (0.06) | **-0.16 (0.15)** | **-0.25 (0.20)** | 0.08 (0.09) | -0.05 (0.05) |
| Flank-to-footprint ratio |  | 0.04 (0.07) | **0.37 (0.13)** | **0.13 (0.15)** | -0.04 (0.07) | 0.04 (0.05) |
| Motif score |  |  | 0.06 (0.08) | 0.06 (0.06) | -0.03 (0.04) | 0.03 (0.04) |
| DNase reads |  |  |  | **0.45 (0.14)** | **-0.19 (0.12)** | 0.07 (0.08) |
| DHS flag |  |  |  |  | **-0.24 (0.13)** | **0.10 (0.08)** |
| Dist. nearest TSS |  |  |  |  |  | **-0.12 (0.08)** |

Second, we took each feature out of the full LR-DNase model separately and compared the performance of the reduced model in comparison with that of the full model (<u>**Figure 4**; **bottom**</u> panel). On average, removing *DNase reads* led to the largest decrease in performance (median decrease compared with the full model was 14.80% in the within–cell-type setting and 12.37% in the cross–cell-type setting). Removing *motif score* led to the second largest decrease in performance (median decrease 4.84%, cross-cell type 3.89%), followed by *flank-to-footprint ratio* (median decrease 1.71%, cross–cell-type 1.43%).

The two approaches together revealed that *DNase reads* was the single most important feature in the LR-DNase model. It was the most predictive single feature, and removing it impacted the performance of the full model most severely. On the other hand, although *flank-to-footprint ratio* was the second most predictive single feature, removing it only had a modest effect on performance, indicating that it is redundant with some of the remaining features, likely the other three features computed from DNase-seq data. Conversely, although *motif score* was not a predictive single feature, removing it had a large effect, indicating that it made a distinct contribution to the full model. Because we found *DNase reads* and *motif score* to be the two most important features for 34% of the datasets by the leave-one-feature-out evaluation, we constructed a baseline logistic-regression model that consisted of just these two features (LR2). LR-DNase achieved higher AUC-PR50 than LR2 for 80% of the TFs (p-value < 2.2×10^−16^; Wilcoxon signed-rank test), showing that the additional five features in LR-DNase contributed significantly.

TFs for which LR-DNase performed particularly well included ELF1, NRF1, YY1, GABPA, USF2, and ZBTB33, datasets with AUC-PR50 ranging from 0.40 (USF2) to 0.88 (ELF1) and containing 3.9% (YY1) to 23.2% (NRF1) positives. LR-DNase also performed well on BDP1 and BRF1, both subunits of the RNA polymerase III initiation factor TFIIIB. The LR-DNase models for these two TFs substantially outperformed LR2 in the K562 cell type (from 0.43 and 0.24 to 0.80 and 0.71, respectively). This result is in concordance with the average profiles shown in <u>**Supplemental Figure 1**</u>, displaying a clear strand-specific cleavage increase in the flanking regions. Moreover, the contributions of the *flank-to-footprint ratio* feature to the full model were 16% and 14% for BDP1 and BRF1, respectively.

**Figure 4.**
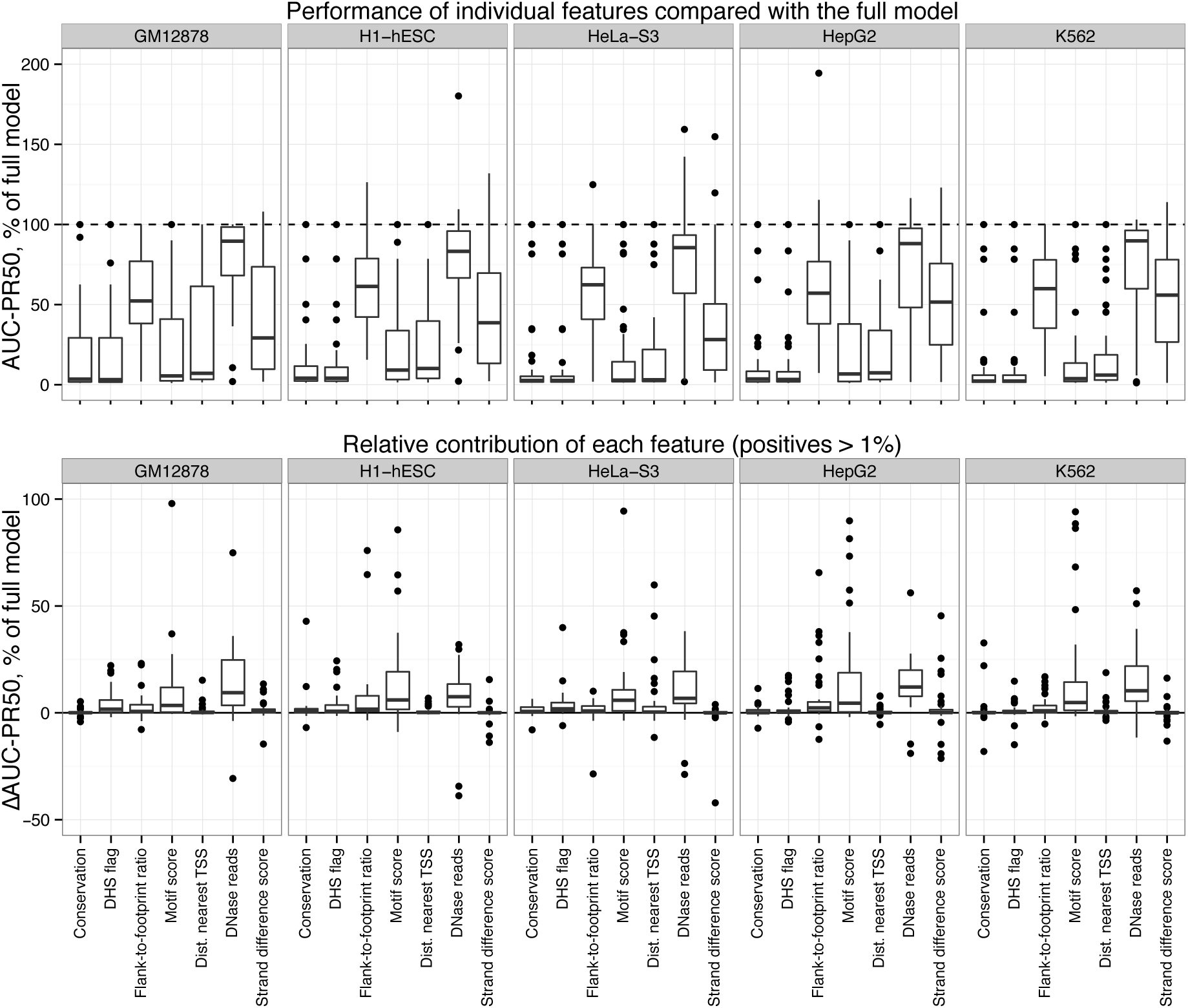
Performance of each feature in isolation and relative contribution of each feature to the full LR-DNase model. (Top) AUC-PR50 of individual features as the percentage of the AUC-PR50 of the full LR-DNase model with all seven features. (Bottom) Relative contribution of each feature measured as the difference in AUC-PR50 upon leaving out the feature of interest, as a percentage of the performance of full LR-DNase model. The outliers were mainly due to the division by small numbers (the pseudocount for computing the divisions was set to 0.01). For clarity not all outliers in the top panel are shown; see Supplemental Figure 11 for the full figure will all outliers.

We performed a pairwise all-to-all statistical comparison of the performance of LR-DNase, its leave-one-feature-out models and single feature rankings, arranged in a directed graph to illustrate their relative performance (<u>**Figure 5**</u>). In this graph the comparisons with the single-feature rankings are in purple and the leave-one-feature-out models are in orange. We also include in this graph four previously published methods in black, which we will discuss three subsections later. Each edge in the graph indicates that the method in the originating node had significantly higher AUC-PR50 than the method in the destination node, based on within–cell-type comparisons. The width of the edge is proportional to the −log_10_(p-value) of the Wilcoxon signed rank test used to compare the two methods, after Bonferroni multiple testing correction. Redundant edges were removed from the graph. This statistical analysis confirmed that LR-DNase performed significantly better than most of the leave-one-feature-out models and single feature rankings, as described above and indicated by outward edges from LR-DNase.

**Figure 5.**
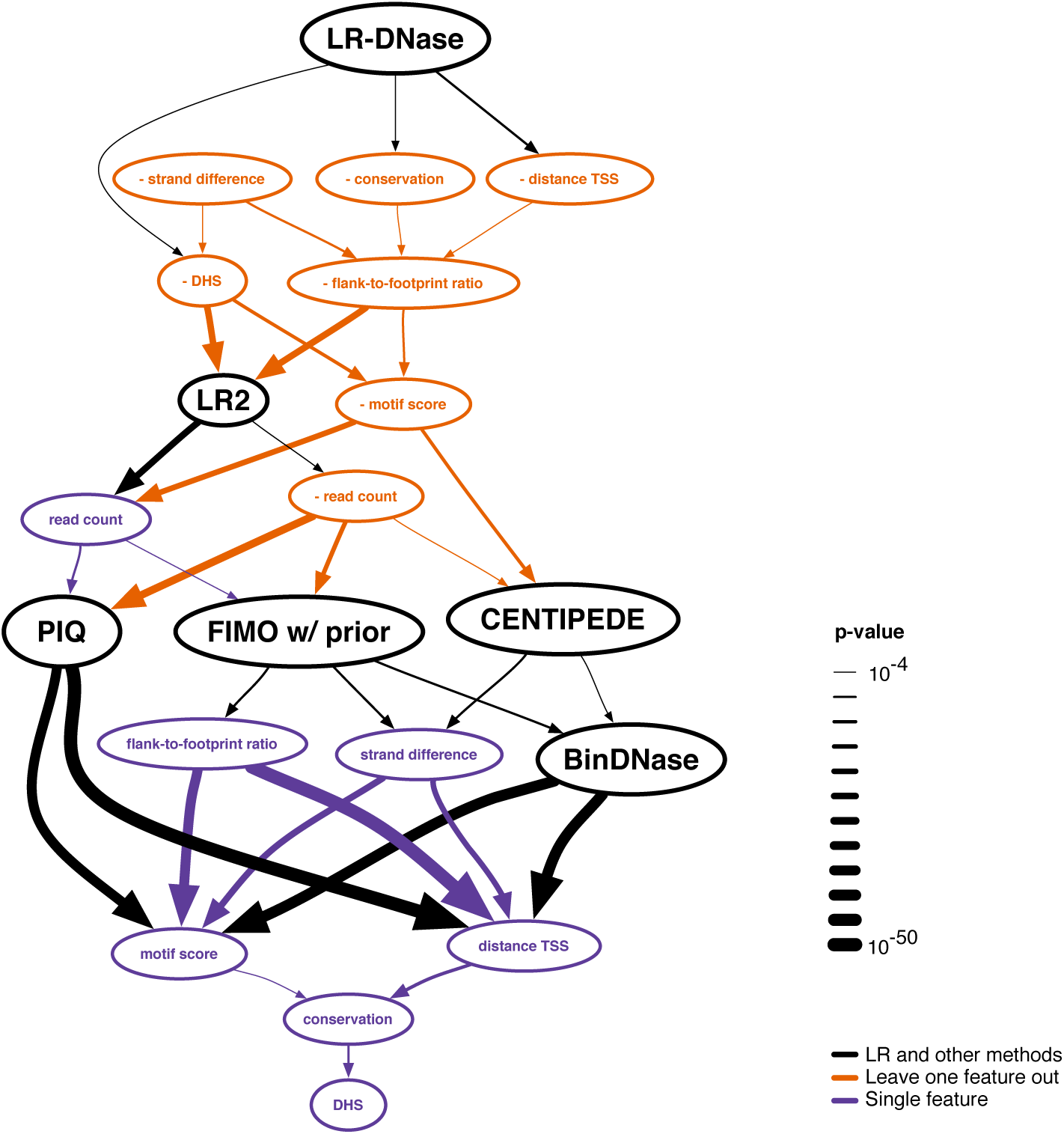
A directed graph comparing the performance of all tested models and methods. Each node represents a model or a method and edges represent significant improvement over the node pointed to. The linewidth of an edge is proportional to Bonferroni corrected −log(p-value) based on Wilcoxon signed-rank test. Only non-redundant edges are shown. Main methods are in black, LR-DNase models with one feature left out are in orange, and single feature rankings are in purple.

The model in which only the *DNase reads* feature was removed performed slightly worse than LR2 which contained just *DNase reads* and *motif score*, indicating the importance of *DNase reads*. Cuellar-Partida et al.(17) also showed that ranking by DNase reads alone outperformed existing methods on a site-centric benchmark. Furthermore, Gusmao et al. showed that ranking the output of existing methods by DNase reads (within the predicted classes) increased their performance (36).

We did not find a significant difference between LR-DNase and the model with the *strand difference score* removed, because they perform similarly across all datasets (<u>**Supplemental Figure 4**</u>), suggesting that *strand difference score* is redundant with the other six features in LR-DNase. We found that the difference in performance between LR2 and the full model from which the *motif score* feature was removed varied widely across the datasets; consequently, there is no edge between the two models in the graph (p-value=0.97, Wilcoxon signed rank test, <u>**Supplemental Figure 5**</u>). The *DHS flag* single-feature ranking is at the bottom of the graph, because it is a binary feature that, when ranked, results in mostly ties.

### Feature contributions are TF specific

Because DNase cleavage profiles are TF specific (8–10), we hypothesized that the relative importance of the LR-DNase features might also be TF specific. First, we performed hierarchical clustering of the LR-DNase models of the TFs based on the weights assigned to the features (heat map in <u>**Supplemental Figure 6)**</u>. As expected, models for the same TFs in different cell types tended to cluster together, which supports the abilities of these models to perform cross– cell-type predictions. Similarly, the models for interacting TFs also clustered (e.g., CTCF, SMC3, and RAD21; JUN and FOS; and MYC, MAX, and MXI1). A number of TFs that tend to bind at promoters (e.g., TAF1, E2F1, MYC/MAX and ZNF143) formed a cluster at the bottom of the heat map, with large and negative weights for *Distance to nearest TSS*. The models for ZNF274 distinctively assigned high weights to *evolutionary conservation score*, *motif score* and *distance to the nearest TSS*, because ZNF274 is a repressive TF that binds at the 3´-end of genes encoding Zinc-finger TFs (37). Also, repressive TFs clustered together near the bottom of the heat map, showing large weights for *Motif score*: REST, MAFK, MEFF, ZNF274. For some of the models for BDP1 and BRF1, the weight for *flank-to-footprint-ratio* was negative (e.g., the datasets with accessions ENCFF002CVP and ENCFF002CVR), reflecting the distinct peak centered at the motif site bound by the TFs.

As a second approach, we looked at the relative impact of each feature on the AUC-PR50. This analysis suggested that the most important features were *DNase reads*, *motif score*, and *distance to the nearest TSS* for 47%, 27%, and 10% of the datasets, respectively. For 34% of the datasets, the two most important features were *DNase reads* and *motif score*. <u>**Figure 6**</u> illustrates the feature contributions for seven example TFs, measured by the decrease of AUCPR50 after removing each feature (<u>**Supplemental Figure 7**</u> for all TFs). USF1 and USF2 are “typical” TFs: they recognize the same motif, and *DNase reads* was the most important feature for both of them. Nevertheless, for USF2 *flank-to-footprint ratio* was the second most important feature, while *motif score* was less important, and the reverse was true for USF1. *DNase reads* and *motif score* were equally important for NRF1. For GABPA, *DNase reads* and the *distance to the nearest TSS* showed similar levels of performance. REST and CTCF are examples for “atypical TFs”. As revealed in the above analysis of feature weights, *Motif score* was the most important feature for repressive TFs such as REST, MAFK, MAFF, and ZNF274, and removing it led to a 95.35% decrease in the AUC-PR50 of ZNF274. CTCF and its paralog CTCFL showed drastically different feature contributions, as described in the next section.

**Figure 6.**
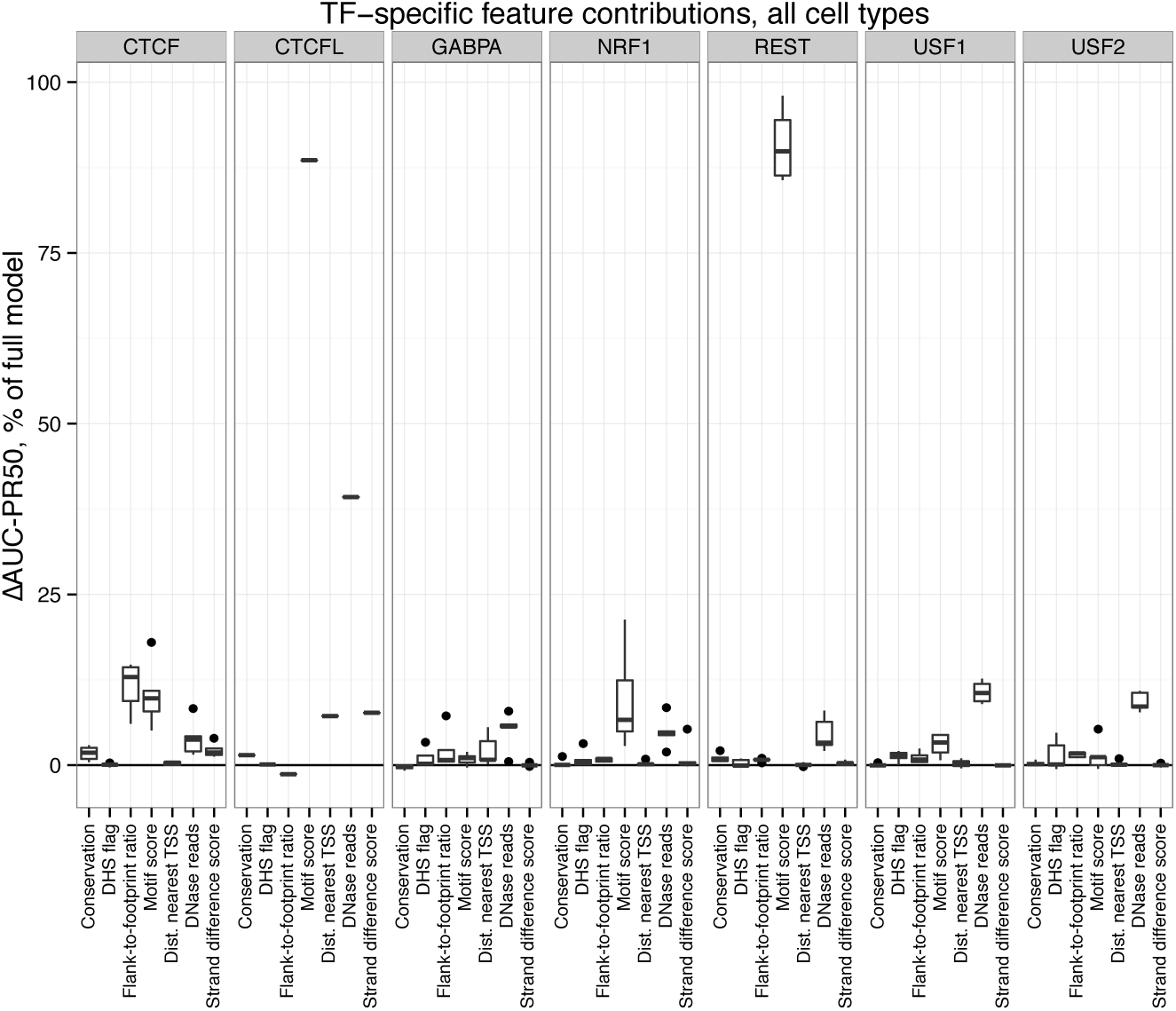
Relative feature contributions for specific TFs, measured as the difference in AUC-PR50 upon leaving out the feature of interest, as a percentage of the performance of full LR-DNase model. Different features contribute the most for different TFs, even for TFs that recognize the same motif.

### Different features were important for predicting CTCF and CTCFL binding

The insulator binding protein CTCF and its ortholog CTCFL (also named BORIS) share a conserved zinc-finger DNA binding domain and, as a consequence, recognize the same motif (22, 38). CTCF is expressed ubiquitously in all cell types, whereas CTCFL is exclusively expressed in testis and is aberrantly expressed in a wide range of cancers (39). The ChIP-seq dataset for CTCFL that we analyzed was collected on K562 cells; thus, we compared it with the CTCF ChIP-seq dataset on K562 cells. Of the 127,309 potential binding sites that were detected using FIMO with the CTCF position-weight matrix, 8,194 were bound by CTCFL, much fewer than CTCF (28,520 sites), and 7,591 sites were shared. Thus, there were many motif sites that were bound by both CTCF and CTCFL or just bound by CTCF, producing an average cleavage profile that was not predictive of CTCFL-specific binding.

*Flank-to-footprint ratio* and *motif score* were the two most important features for predicting CTCF binding in K562 cells, and *distance to the nearest TSS* was unimportant. In contrast, *motif score* was by far the most important feature for predicting CTCFL binding, and *DNase reads* was the second most important feature, followed by *distance to the nearest TSS* (<u>**Figure 6**</u>). We examined in detail the differential importance of these features for the two TFs.

We first investigated why *flank-to-footprint ratio* was less important for CTCFL. <u>**Supplemental Figure 8**</u> shows the average DNase cleavage profiles for sites bound exclusively by either CTCF or by CTCFL. We observed a less pronounced profile at sites exclusively bound by CTCFL than at sites bound by CTCF, although a higher overall cleavage rate throughout the entire ±200bp region centered on CTCFL binding sites. Additionally, the peak at the +4 position, which is the signature of CTCF binding sites, is absent in the CTCFL profile, hence the lower importance of the *flank-to-footprint ratio* feature for CTCFL.

We then examined why *distance to the nearest TSS* was important only for predicting CTCFL binding. The median distance from sites bound exclusively by CTCFL to the nearest TSS was much shorter than the distance for the sites bound exclusively by CTCF or sites bound by both TFs (539 vs. 15,350 bp and 6,968 bp; p-value < 2.2×10^−16^ by Wilcoxon signed-rank test).

Finally, we investigated why *motif score* was particularly important for predicting CTCFL binding. Pugacheva et al. found that CTCFL preferentially binds at genomic regions that contain clusters of CTCF motif sites and that many of these regions were active promoters and enhancers, often bound by CTCF as well (38). However, CTCFL-only peaks did not have more motif sites than CTCF-only peaks and we found that the motif score for CTCF-only peaks were significantly higher than for CTCFL-only peaks (p-value = 1.26 × 10^−11^, Wilcoxon test, <u>**Supplemental Figure 9**</u>).

### Comparison of LR-DNase and LR2 with other state-of-the-art methods

We compared LR-DNase with four previously published, state-of-the-art methods: three unsupervised methods (CENTIPEDE, PIQ, FIMO with epigenetic priors) and one supervised method (BinDNase)(17–20). Statistical comparison of the results (<u>**Figure 5**</u>) shows that all four methods underperformed LR-DNase and its leave-one-feature-out models as well as the baseline model LR2. The performance of PIQ, CENTIPEDE, and FIMO with epigenetic prior were statistically indistinguishable from one another, indicated by the lack of edges among them in the graph, while FIMO with prior and CENTIPEDE significantly outperformed BinDNase.

For a direct comparison with LR2, we evaluated each method on the 243 non-redundant datasets, calculating the change in AUC-PR50 as a fraction of the maximum possible improvement (FMI) upon LR2 (see Methods). <u>**Figure 7**</u> shows the relative performance, evaluated using FMI, of all tested methods, models and single feature rankings. We observed a clear difference between previously published methods and LR-DNase. Overall, LR-DNase and LR2 outperformed CENTIPEDE, PIQ, FIMO with epigenetic priors, and BinDNase. For example, CENTIPEDE showed a decrease of 28.00±38.51% (median 11.26%) of that of LR2.

**Figure 7.**
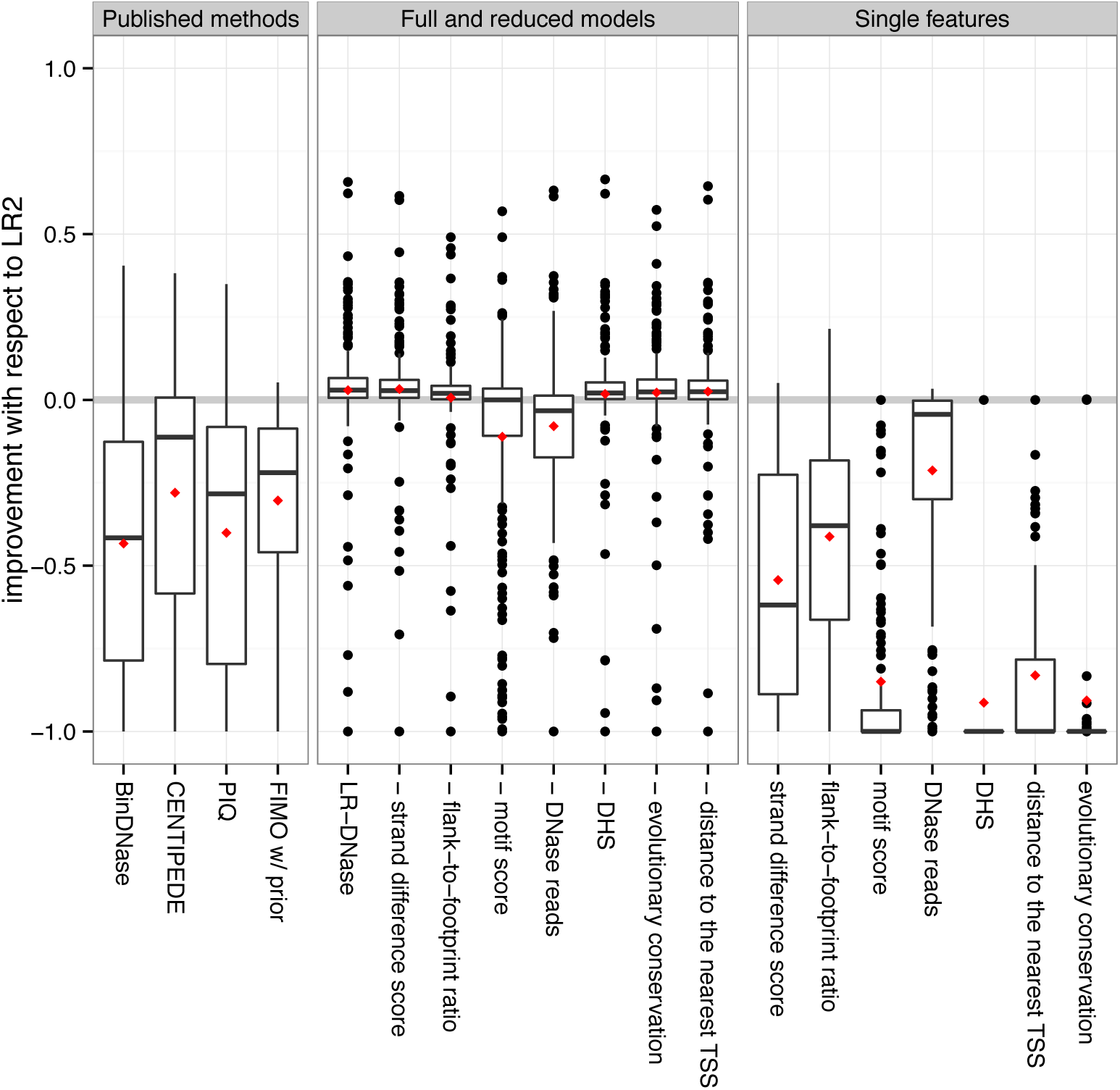
Boxplots of the fraction of maximum possible performance improvement gained by each of the LR-DNase models and four published methods BinDNase, CENTIPEDE, PIQ and FIMO with epigenetic priors as compared with the baseline model LR2 (see Methods). Means are indicated as red diamonds.

For a small set of TFs, including CTCF and its cofactors SMC3 and RAD21, as well as ZBTB33 and USF2, all methods performed well. Interestingly, ELF1 and GABPA were predicted well by most methods except CENTIPEDE (<u>**Supplemental Figure 10**</u>). For these two TFs the *distance to the nearest TSS* was highly informative; ranking solely by that feature yielded AUC-PR50 values of 0.52±0.10. However, the performance of LR-DNase for these two TFs did not change significantly when the *TSS distance* feature was removed; this may be because the *DNase reads* and *TSS distance* features were correlated for these TFs (r=–0.45±0.05). *Distance to the nearest TSS distance* was also a feature for CENTIPEDE, and it is not known why CENTIPEDE did not perform well for these TFs.

In general, the different methods performed quite similarly to one another across TFs. We observed a Spearman correlation of 0.97 (p-value < 2.2x10^−16^) between the AUC-PR50 values of the LR-DNase model and the AUC-PR50 values obtained by taking the highest performing method among the four previously published methods for each dataset (<u>**Supplemental Figure 11**</u>). Thus, LR-DNase was not particularly effective on the subset of TFs for which the other three methods performed poorly.

### Performance on TFs that prefer to bind outside DHS

Most TFs prefer binding within DHSs (<u>**Supplemental Table 2**</u>); nevertheless, for a minority of TFs (ZNF274, MAFK, MAFF, REST, BRF2, CEBPB, USF1, SPI1 and FOXA1, in 26 out of the 243 non-redundant datasets), over 50% of the bound sites are outside DHSs. Because only 3-5% of the nucleotides in the genome are in DHSs, even for these TFs (with the exception of ZNF274), their binding sites are still enriched in DHSs. Among these TFs, ZNF274, MAFK, MAFF, and REST are repressors (37, 40, 41). FOXA1 is a pioneering TF which can bind to closed chromatin regions and open up the chromatin for other TFs (42).

Not surprisingly, given that by definition there are fewer DNase-seq reads outside of DHSs, we found that for these TFs the most important feature is not *DNase reads* but *motif score* (see <u>**Supplemental Figure 12**</u> and <u>**Supplemental Figure 13**</u>). ZNF274 shows a strong negative coefficient for the *DHS* feature in our model; however, the LR-DNase model showed low performance for this TF (AUC-PR50=0.031±0.052).

## DISCUSSION

The advent of high-throughput DNase-seq assays has greatly enhanced our ability to detect regulatory elements and TF binding sites, and can potentially determine genome-wide occupancies of all TFs at once. A number of algorithms have been developed to predict TF binding sites using DNase-seq data, and they generally fall into two types. One type of method, such as CENTIPEDE, FIMO with priors, PIQ, and BinDNase (20), uses the knowledge of the sequence motifs available for many TFs to scan the genome and find all motif sites, and then predicts which of these sites are bound by the TFs. The other type of method, such as Wellington, HINT and DNase2TF (11–13), performs *de novo* predictions using just the cleavage pattern of DNase I, segmenting the genome into short regions that are bound by a TF (without specifying which TF) and regions that are not. The former type of method is better able to control false positives, especially for TFs with high sequence specificities, but only the latter type of method can discover the binding sites of novel TFs. We have introduced a new method, LR-DNase, that belongs in the former category, and we compared it with other methods of this type.

A gold standard is required for evaluating the performance of LR-DNase and comparing it with other methods. A widely used gold standard is ChIP-seq data of specific TFs in the same cell type as the DNase-seq data; however, a major difficulty in constructing the gold standard is the difference in resolution between ChIP-seq data and TF binding. ChIP-seq peaks for TFs are typically ~250 bp long, which is a much lower resolution than the individual binding sites of TFs, 6-25 bp long. In order to construct a gold standard at base-pair resolution from the lower resolution ChIP-seq data, two approaches are commonly used: site-centric, which restricts the analysis to only motif sites that score higher than a lenient cutoff, denoting the sites inside ChIP-seq peaks as positives and outside as negatives; and peak-centric, which picks the best scoring motif site in each ChIP-seq peak as the positive while treating all other positions in the genome as negatives. Both approaches have limitations, and we opted for the site-centric approach because our LR-DNase algorithm is a classifier that uses features computed for individual motif sites. These features are ill defined for genomic positions that are not motif sites. In addition, the site-centric evaluation more accurately reflects direct TF binding, as opposed to indirect binding via the interaction of different TF proteins, since the direct binding sites are likely to pass a minimal motif cutoff. Nevertheless, a substantial proportion of ChIP-seq peaks do not have any motif sites that pass our selected motif score cutoff. These peaks could be the result of indirect binding, which is particularly likely in the case when the peaks are enriched for the binding sites of another TF. Alternatively, these peaks could also be due to noise in the ChIP-seq experiment, or could be due to DNA shape or higher-order effects that enable a piece of DNA that does not contain a classically defined PWM motif site to nonetheless be bound by the TF. We argue that these scenarios would place challenges equally for LR-DNase and other algorithms, and hence the comparison would still be fair using the site-centric approach. Indeed, the site-centric evaluation was performed for the other algorithms CENTIPEDE, PIQ, and BinDNase (20), and FIMO with priors was evaluated using both site-centric and peak-centric methods.

Based on site-centric evaluation, LR-DNase performed well above the null model and significantly better than four other methods, CENTIPEDE, PIQ, BinDNase, and FIMO with priors. LR-DNase achieved higher AUC-PR on datasets with higher percentages of true positives, whereas datasets with <1% positives posed major challenges. Neither LR-DNase nor the other methods could predict at the false positive rate of 5%—at this FDR rate, LR-DNase could achieve greater than zero sensitivity for only 50% of the datasets. Thus, these methods need to be combined with downstream methods performing high-throughput validation.

Because LR-DNase is a logistic regression based method, we could perform a detailed evaluation on the relative contributions of its seven features, four of which were derived from DNase-seq data. We found that even though each of these four features was individually predictive, there was substantial correlation among them. Consequently, *DNase reads* was the most predictive feature, and the other three features provided only moderate increases in predictive power. We also found that *motif score*, although not a highly predictive feature by itself in the site-centric evaluation setting, substantially augmented the predictive power of *DNase reads*. As a result, a logistic regression model based on just the two features, *DNase reads* and *motif score*, performed only moderately worse than the full LR-DNase yet better than CENTIPEDE, PIQ, BinDNase, and FIMO with priors. Earlier papers also indicated that DNaseseq reads alone often outperformed state-of-the-art prediction methods (17, 36). These two features do not capture the high-resolution “footprints” left on DNase cleavage profiles by bound TFs; thus, currently no method has fully harnessed the information in ultra-deep DNase-seq datasets.

There is great interest in the use of ultra-deep DNase-seq data for TF binding site detection, and there is also an ongoing discussion on the limitations of digital footprinting (43, 44), in particular for TFs that have short-lived interactions with DNA. He et al. and Sung et al. reported that the average binding signal that is usually observed at bound motif sites may be partly the result of sequence-specific cleavage by the DNase I enzyme, suggesting that the shape of the high-resolution DNase cleavage profile might be confounded with information that has not been previously modeled (10, 13). We have shown that low-resolution features, such as *DNase reads* and *flank-to-footprint ratio*, were effective for many datasets. We tried to correct for the sequence-specific cleavage bias by the DNase I enzyme prior to computing the features for LR-DNase, but this did not lead to improved performance (data not shown). One active future direction is to design algorithms that can take advantage of the fine features in DNase cleavage profiles offered by ultra-deep DNase datasets.

For many ChIP-seq datasets, the peaks were not enriched in the canonical binding motif for the interrogated TF but instead contain the canonical motifs of other TFs, indicating the ChIP-seq peaks could be caused by co-binding with or indirect binding between multiple TFs (22). This observation also suggests that methods that identify the motifs of multiple TFs and model their interactions, as well as hybrid methods leveraging both TF-specific sequence information, cleavage data and histone modification data, could provide future avenues for research.

A number of datasets included paralogous TFs, such as CTCF and CTCFL, that share the same sequence motifs. It is a challenge to predict the sites that are bound specifically by a paralogous TF, because the observed DNase cleavage patterns could be produced by its paralogs. As we showed for CTCF and CTCFL, different features contributed the most to predicting their respective binding sites. One future direction is to consider all expressed TFs that share the same motif with detailed modeling of single, co-binding and competitive binding scenarios.

